# Relative value learning in *Drosophila melanogaster* larvae

**DOI:** 10.1101/2025.09.03.673910

**Authors:** Sadniman Rahman, Nobuaki K. Tanaka, Michael Schleyer

## Abstract

The ability to learn from past experiences to inform future decision-making is crucial for humans and animals alike. One question with important implications for adaptive decision-making is whether we learn about the absolute values of cues we encounter (how good or bad?), or about their relative values (how much better or worse than the alternative?). Humans have been shown to use relative value learning, even when it leads to suboptimal decisions. In this study, we ask whether insects employ absolute or relative value learning. Using the larvae of the fruit fly *Drosophila melanogaster*, we designed associative odour-taste learning experiments to distinguish both kinds of learning and find that larvae learn about the relative rather than the absolute values of both rewards and punishments, irrespective of the number and sequence of training trials. This suggests that relative value learning is a facility shared across the animal kingdom from maggots to humans, and can be accomplished even by small insect brains. Given the potential of *D. melanogaster* as a model organism for in-depth neurobiological analyses, our study opens up an opportunity to reveal the detailed mechanisms underlying relative value learning.

## Introduction

One of the most fundamental functions of our memory is to guide adaptive decision-making [1–3]. From what to eat for dinner to which work project to prioritize, decision-making pervades our daily lives. A wealth of evidence suggests that choice options are evaluated in a relative rather than absolute manner, that is, we rate an option according to how it compares to others [4–6]. In many cases, our previous experiences help us to correctly assess the value of each option and to select the best choice. How exactly value is assigned to a given option during such previous experiences is the subject of intensive research.

A common approach to study this question is associative learning. In the last decade, several studies have shown convincingly that humans apply relative evaluation of rewards and punishments during associative learning. This means that humans learn the value of a reward or punishment relative to other rewards or punishments that were available in a particular training context (how much better or worse?) rather than an absolute value (how good or bad?) that would be independent from the training context. This process is known as relative value learning, adaptive value coding, value normalization or range adaptation [7–13]. This discovery is important for our understanding of economic, value-based decisions because relative value learning can lead to suboptimal choices. For example, an option that had been paired with an objectively low value, but in a context of even lower values, is preferred over another option that had been paired with an objectively higher value, but in a context of even higher values. In other words, humans tend to choose options that have been better than the alternatives in previous experiences, even if this implies choosing the objectively worse of the options at hand [8, 14–15].

The last observation raises the question of the adaptive significance of relative value learning. Proposed advantages include a higher efficiency of saving relative value differences rather than absolute values [16–17], and an increased sensitivity to the most likely outcomes in a given environment [18]. Similar proposals have been made in the context of the optimal foraging and decision-making theory [19–20]: in stable environmental conditions in which the availability of different options is known to the forager and does not change over time, learning absolute values of food sources is of advantage and foragers should accept food sources based on a fixed, absolute threshold. In contrast, in fluctuating environments foragers profit from learning constantly updated values relative to their internal state and previously encountered food sources because it provides flexibility, improves cost efficiency and allows to maximize food intake [21–28].

Relative value learning has been found in mammals and birds, suggesting that the principles of value coding are shared across vertebrates [29–35]. Invertebrates like insects have very differently structured brains that are much smaller and often regarded as too simple to allow for complex tasks. Nevertheless, associative learning has been demonstrated in insects like bees, crickets and flies since decades [36–38], and relative evaluation of food sources has been observed during foraging [28]. Several insect species, including flies, bees and ants, have been demonstrated to be capable of discriminating values of different rewards or punishments and form memories of different strength accordingly [39–44]. However, these studies cannot answer the question whether insects actually learn about, that is, “save” the relative value of a reward or punishment, or the absolute value. A study in grasshoppers showed that insects can learn a reward value relative to their current internal state when being trained at different hunger states [45]. Honeybees have been found to anticipate increasing and decreasing rewards, and thus to learn a reward value relative to previous instances of the same reward [46]. Still, the question remains whether insects compare and learn about the relative values of different rewards and punishments that are present during training, even if this leads to suboptimal decisions, like vertebrates do.

Here, we use the larval stage of the fruit fly *Drosophila melanogaster*. In the past decades, the *D. melanogaster* larva has established itself as useful model organism for neurobehavioural research [47–51]. It combines a small brain with only about 10,000 neurons and a rich toolbox for transgenic manipulations of individual cells with a surprisingly complex behavioural repertoire. This includes appetitive and aversive classical as well as operant conditioning, long-term-memories, decision-making and adaptive outcome expectations [39, 52–61].

We designed a behavioural paradigm to distinguish absolute and relative value learning in classical conditioning. Based on established odour-taste associative experiments [52, 62–64], we systematically ask whether larvae learn about the relative values of rewards as well as punishments and whether this learning depends on the odours presented during the training or the memory retrieval test, the number of training trials and the sequence of training trials. Our findings will help to shed light on how insects encode values and provide an opportunity to investigate the neuronal mechanism of relative value learning in a numerical simple brain.

## Materials and Methods

### Animals

Throughout the study, we used 4 days old, third instar, feeding-stage Canton-S wild-type strain of *Drosophila melanogaster* larvae. We maintained them on the standard fly food at 25°C with a 12:12 h light–dark cycle.

### Three trials fructose learning experiment

The associative learning experiments used in this study are based on a commonly used experimental paradigm [63], with some deviations. We trained larvae with two odours, *n*-amylacetate (AM; CAS: 628-63-7, Tokyo Chemical Industry Co., Ltd, Japan) and 1-octanol (OCT; CAS: 111-87-5, Tokyo Chemical Industry) diluted 1:49 and 1:1 in paraffin oil (CAS: 128-04375, Fujifilm Wako Pure Chemical Corporation, Japan), respectively, associated with different concentrations of fructose (CAS: 127-02765, Fujifilm Wako Pure Chemical). Specifically, we used 0.1 M (+) and 2 M (++) mixed in 2 % agar (CAS: 016-11875, Fujifilm Wako Pure Chemical) as weakly and strongly rewarding substrates, respectively, as well as pure, tasteless agar as neutral situation (∅). Every cohort of larvae was trained with each odour being paired with one of the three different substrates. A total of nine combinations were tested in this experiment, summarized in Table 1.

**Table 1:** Experimental design. AM: *n*-amylacetate, OCT:1-octanol, ∅: neutral substrate, +: weak reward, ++: strong reward. The group numbers in brackets are referred to in the text.

|  |  | Reward paired with AM |  |  |
| --- | --- | --- | --- | --- |
|  |  | Ø | + | ++ |
| Reward paired with OCT | Ø | AMØ/OCTØ (1) | AM+/OCTØ (4) | AM++/OCTØ (7) |
|  | + | AMØ/OCT+ (2) | AM+/OCT+ (5) | AM++/OCT+ (8) |
|  | ++ | AMØ/OCT++ (3) | AM+/OCT++ (6) | AM++/OCT++ (9) |

At first, approximately 20 larvae were taken from the food vials, gently washed with tap water and placed at the centre of a training Petri dish (9 cm inner diameter, order #3-1491-01, AS ONE, Japan), filled with one of the three substrates. Two custom-made teflon odour containers filled with the same odour (either AM or OCT) had been placed on opposing sides of the dish just before start of the experiment. The larvae were allowed to move freely over the dish for 2.5 minutes. Subsequently, they were gently transferred to another training dish, filled with one of the three substrates and with the respective other odour placed, and stayed there for another 2.5 min. We repeated this cycle for 3 times. For example, to train group 6 in Table 1 (AM+/OCT++; Fig. S1a), the larvae were placed in a training dish featuring two odour containers filled with AM and the weak reward (AM+), followed by a second dish featuring OCT and the strong reward (OCT++). In half of the repetitions of the experiment, the sequence was reversed such that the training started with OCT++ followed by AM+. After three training cycles, we immediately tested the animals’ preference for AM by placing them at the centre of the test Petri dish which featured on one side an odour container filled with AM and on the opposite side an empty container (EM). All experiments in this study used test dishes filled with 5 mM quinine, unless explicitly mentioned otherwise (for more details, see below). After 3 min, the number of larvae (#) on the AM side, on the EM side, and in a 10 mm wide middle zone was counted to calculate their preference for AM using the following formula. Larvae on the lid were excluded from the analysis.

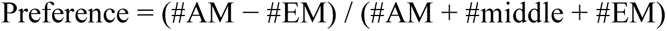

Thus, the preference is limited between −1 and +1, with negative scores indicating avoidance and positive scores indicating approach to AM. In some experiments, we calculated a memory score from the preferences of two reciprocally trained groups, following the established procedure [63]. In these cases, always the preference of a group in which OCT had been rewarded and AM not (e.g. AM∅/OCT++) was subtracted from a group with opposite contingency (AM++/OCT∅), divided by two:

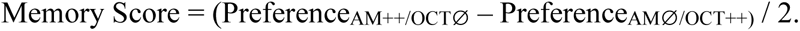

Thus, the memory score is limited between −1 and +1, with negative scores indicating aversive, and positive scores indicating appetitive associative memory.

For further statistical analyses, we estimated the subjective value (SV) of each of the three substrates from the preference behaviour of the animals. Towards this end, we used the mean preferences of groups 1, 4 and 7 (see Table 1), as in each of these groups AM was paired with a different substrate while OCT was paired with the neutral substrate. We reasoned that these groups are best suited to estimate the value the animals assign to each substrate, independent of whether they learned the absolute or relative value. Specifically, group 1 (AM∅/OCT∅) did not involve any rewards and thus arguably no associative learning and was therefore used to determine the animals’ baseline AM preference after the training procedure but in absence of an associative memory. Groups 4 (AM+/OCT∅) and 7 (AM++/OCT∅) were then used to estimate the SV of the weak and strong reward, respectively. We calculated the SV of each substrate as follows:

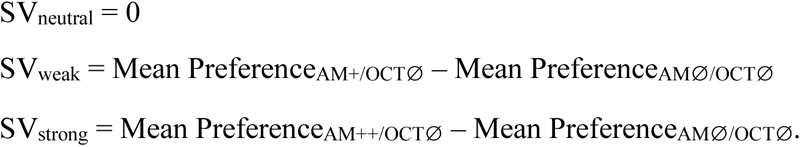

From these SV values, we then calculated the difference in value between the two training substrates:

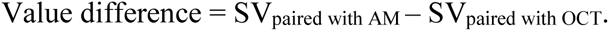

### Variations of the fructose learning experiment

We performed some variations of the experiment described above, always following the same experimental design with 9 combinations of neutral substrate, weak and strong reward, and always testing for the preference of one odour as described.

- Testing for OCT: in one experiment, the animals were trained exactly as described, but the preference for OCT was measured by presenting OCT instead of AM in the test Petri dish. The preference, SV and value difference were calculated analogously for OCT instead of AM.
- Training with one odour: the experiment was performed while using only one odour during training, replacing the other odour with empty containers (Fig. S1b). Such one-odour training is well established [62]. We used either AM, OCT or undiluted Benzaldehyde (BA; CAS: 025-12206, Fujifilm Wako Pure Chemical). For example, instead of training the animals AM+/OCT++ (group 6 in Table 1), they were trained AM+/EM++. The preference, SV and value difference were calculated analogously for the previously trained odour.
- Training with only one training cycle: the training followed the design in Table 1, but each substrate-odour pairing was only presented a single time before testing for the AM preference in an immediate test. This allowed us to compare two possible sequences: either OCT was presented first (e.g. OCT++ → AM+; Fig. S1c) or AM was presented first (AM+ → OCT++; Fig. S1c’). In an additional experiment, we omitted both odours during the training phase, merely exposing the animals to different sugar concentrations before testing them for their AM preference.

### Quinine learning experiment

This experiment was performed in the same way, except using two concentrations of quinine (CAS: 177-00462, Fujifilm Wako Pure Chemical), 0.1 mM (-) and 5 mM (--) as weakly and strongly punishing situations (Table 2). In an analogous experiment, we used 9 different combinations of a neutral substrate (pure agar), reward (2 M fructose) and punishment (5 mM quinine). This included groups of larvae that were trained with one odour being paired with a reward and the other one paired with a punishment (Table 3). Here, we used groups 5-6 to calculate the SV.

**Table 2:**
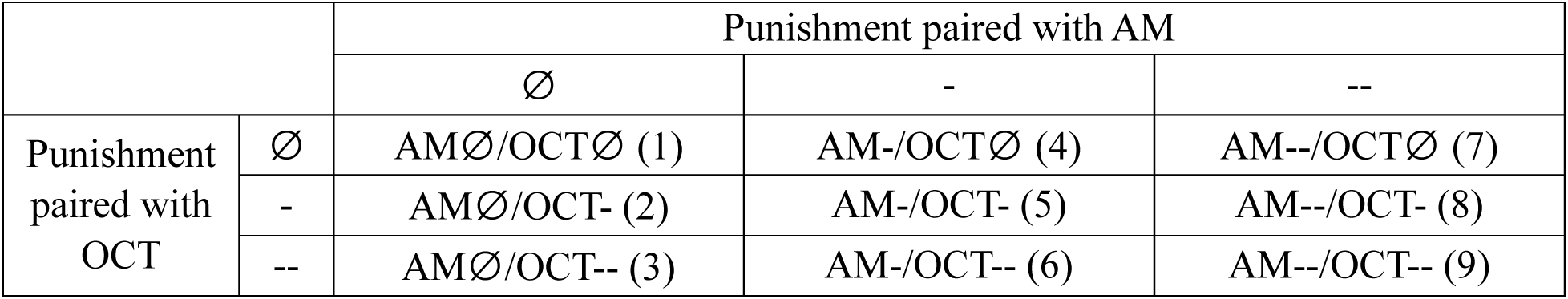
Experimental design of the quinine only experiment. For more details, see Table 1.

**Table 3:** Experimental design of the fructose-quinine experiment. For more details, see Table 1.

|  |  | Substrate paired with AM |  |  |
| --- | --- | --- | --- | --- |
|  |  | -- | Ø | ++ |
| Substrate paired with OCT | -- | AM--/OCT-- (1) | AMØ/OCT-- (4) | AM++/OCT-- (7) |
|  | Ø | AM--/OCTØ (2) | AMØ/OCTØ (5) | AM++/OCTØ (8) |
|  | ++ | AM--/OCT++ (3) | AMØ/OCT++ (6) | AM++/OCT++ (9) |

Previous studies showed that quinine memory is only expressed in presence of quinine, while the expression of fructose memories is independent of the absence or presence of quinine [53]. Therefore, and to maintain comparability across experiments, all experiments in this study used test Petri dishes filled with 5 mM quinine, unless explicitly mentioned otherwise.

### Taste preference experiment

In the learning experiments, the larvae are often exposed to highly concentrated tastants (fructose or quinine) followed by a much lower concentration of the same taste. It is conceivable that after being exposed to a high concentration, the animals’ sensing and/or motivation towards the lower concentration is impaired. To control for this, we placed groups of 20 larvae on a Petri dish with either pure agar or agar containing the highly concentrated tastants (2 M fructose or 5 mM quinine) for 2.5 minutes. Afterwards, the larvae were transferred to a split Petri dish with pure agar on one side and the low concentration of the same tastants (0.1 M fructose or 0.1 mM quinine) on the other side. After 3 minutes, the number of the animals on either side and in a 10 mm middle zone were counted and the taste preference was calculated:

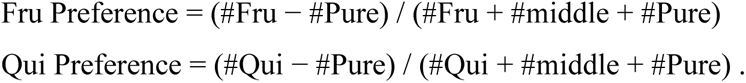

### Statistical analyses

Data are displayed as raw data points, overlaid with the mean as middle line and the 95 % confidence interval as error bars. For all statistical tests described below, when multiple comparisons were performed within one analysis, a Bonferroni-Holm correction was applied to keep the experiment-wide error rate below 5 % [65].

To determine whether gradual increases of reward or punishment strength consistently affects odour preferences and memory scores, Spearman Rank Correlations (SRC) were performed, followed by pairwise Mann-Whitney U tests (MWU) if the SRC was significant. SRC were chosen over Pearson correlations as SRC does not assume a linear relationship of the data, only a monotonically increasing or decreasing relationship. Spearman’s r_S_ values are displayed within the figures to allow readers to judge the strength of the correlations (+1 and −1 indicating perfect positive or negative correlations, respectively). In experiments measuring taste preference, we compared groups with MWU tests and determined whether the preferences were significantly different from zero using the Wilcoxon Signed-Rank test (WSR).Graphs, MWU, WSR and SRC tests were made in Statistica 11 (StatSoft, Tulsa, USA).

As the preference scores in our experiments are limited between −1 and 1, their relationship with any experimental parameter is likely to follow a sigmoid curve. Therefore, we fitted a simplified generalized logistic function to the data. The generalized logistic function can be changed by a number of parameters, of which we used three in this study [66]. The function followed the form:

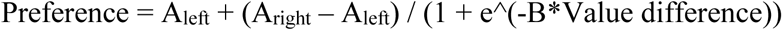

Here, A_left_ and A_right_ indicate the left and right horizontal asymptotes, respectively, and B the growth rate, that is, the steepness of the function. The parameters of the function for each experiment were determined by nonlinear least square fitting using the “nls” function in R [67], and are reported in Supplementary table S1 along with the raw data. To compare the generalized logistic function with a simpler linear function, we compared the Akaike information criterion (AIC) that balances the likelihood of a fitted model with its complexity. The AIC is regularly used to compare statistical models for the same data set, with the model with the lowest AIC being considered the best fit [68].

## Results

### Larvae learn about the relative value of rewards

We performed an associative learning experiment that was modified from the most commonly used learning paradigm in *D. melanogaster* [63, 69]. We trained larvae by alternatingly pairing *n*-amylacetate (AM) with a given concentration of fructose reward and 1-octanol (OCT) without reward, and subsequently tested them for their preference of AM alone (Fig. 1a). The presence of only AM during the test, different from the standard procedure, allowed us to assess what the animals had learned specifically about this odour. Two fructose concentrations were chosen based on initial experiments (Fig. S2a-c). We ensured that the larvae can still detect and process the lower of the two fructose concentrations immediately after being exposed to the higher concentration, although their preference was slightly reduced (Fig. S2d).

**Figure 1.**
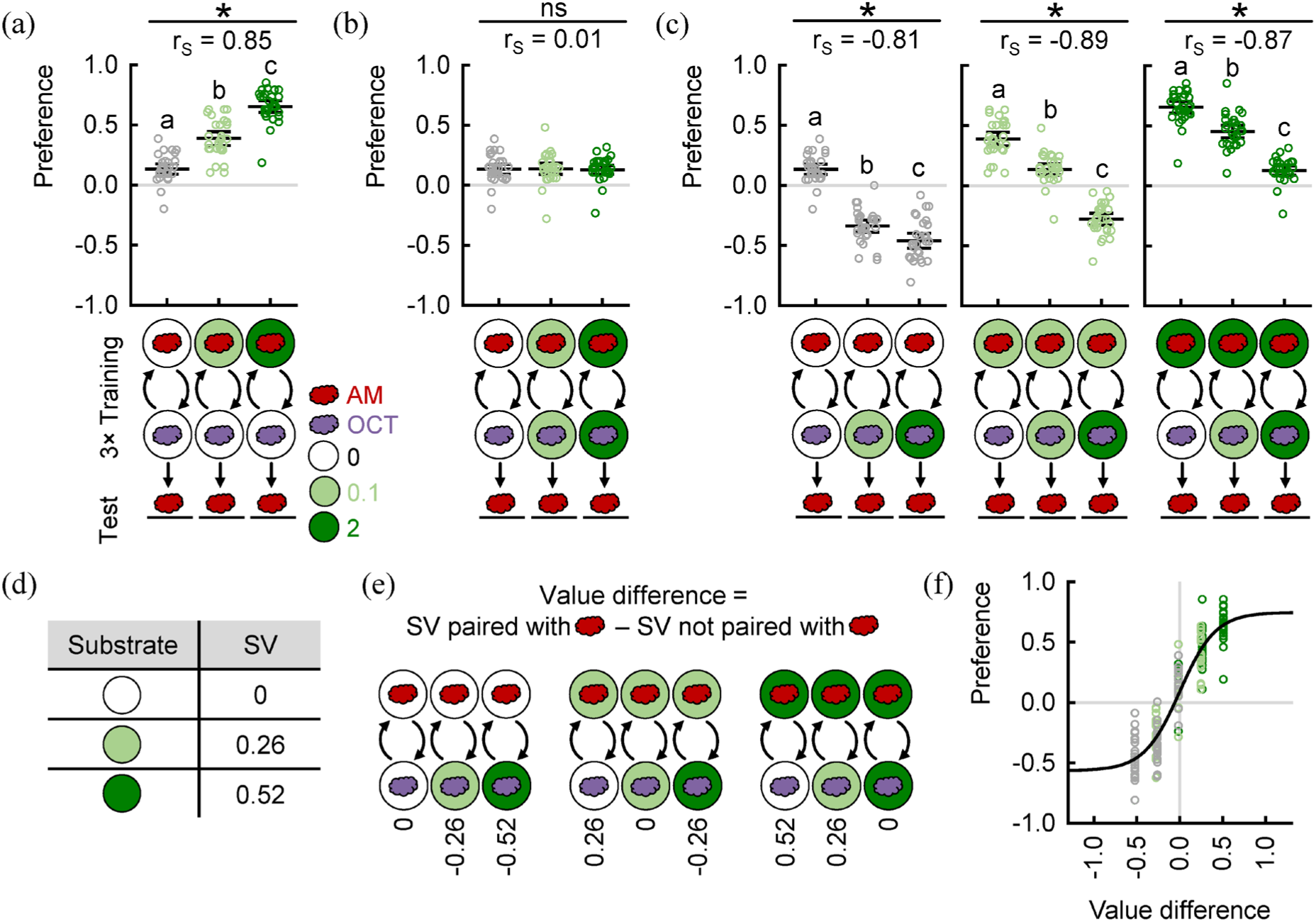
Larvae learn about the relative value of rewards. (a) Larvae were trained with AM paired with 0, 0.1 or 2 M fructose reward and OCT paired with no reward. (b) Larvae were trained with the same fructose concentration in both training dishes. (c) Larvae were trained with the fructose concentration paired with AM kept constant, but the concentration paired with OCT varied. (d) The subjective value (SV) of each reward was deduced from the animals’ preference in (a). See Materials & Methods for details. (e) The (signed) difference in reward value for each experimental group. (f) AM preferences increased with increasing difference in reward value. A generalized logistic function was fitted to the data. Sample size is 30 each. Note that the groups displayed in (a-b) are also included in (c). * and “ns” indicate significant and not significant SRC tests, respectively, and r_S_ indicates the Spearman’s correlation coefficient. Different small letters above each group indicate significant pairwise difference from each other (Mann-Whitney U tests). For all raw data along with the detailed results of the statistical tests, see Supplementary table S1.

Larvae have a positive innate preference to all odours used in this study that can be modulated by non-associative processes [62, 64]. We therefore included a baseline group with the same handling and odour exposure as all other groups, but without any rewards and therefore arguably without associative learning (Fig. 1a, left-most group). Compared to this baseline, we found that the animals’ preference for AM increased with increasing reward concentration, suggesting stronger associative learning (Fig. 1a, see also Fig. S2a-c).

Importantly, in this experiment the larvae could learn to associate AM either with an absolute value of the paired reward (how good it was), or with its relative value (how much better it was than the alternative). As the alternative during training was plain, neutral agar, both hypotheses predict the same outcome and the current experiment does not allow to differentiate between them.

Therefore, we trained the larvae with AM and OCT being paired with the same fructose concentration (Fig. 1b). If the animals learned about the absolute reward value, the substrate that was paired with OCT should not matter and the animals should show the same level of associative learning as in Fig. 1a. If the animals learned about the relative reward value, however, there should be no sign of associative learning in this experiment, because both alternating substrates were the same and therefore the relative value of the reward was zero (neither better nor worse than the alternative). Indeed, we found that the animals all displayed the same baseline AM preference, irrespective of the reward strength (Fig. 1b). This suggests that they learned about the relative and not the absolute reward value.

Next, we modified the experiment such that AM was paired with no reward and thus the absolute reward value was zero, but the alternative featured different reward concentrations (Fig. 1c, left). We found that the AM preference was decreasing with increasing reward strength that had been paired with OCT. Again, this fits to the hypothesis of relative value learning: as the substrate paired with AM had been less good than the alternative, its relative value was negative. Finally, we repeated the same experiment, but paired AM either with the weak reward (Fig. 1c, middle) or the strong reward (Fig. 1c, right). In each case, the AM preference declined along with the reward strength that had been paired with OCT, and in each case the result indicated that the animals’ behaviour was determined by the relative reward value associated with AM. Strikingly, this even caused the larvae to avoid AM after it had been paired with a reward, but the alternative substrate had been even better (Fig. 1c, 6^th^ group from the left). Thus, we concluded that *D. melanogaster* larvae learn about the relative value of a reward, not the absolute value. We obtained an analogous result when we repeated the complete experiment but tested for the OCT preference instead of AM (Fig. S3a-c).

In this study we used both fructose reward and quinine punishment (see below). Quinine memories have been repeatedly shown to be only retrieved when the animals are tested in presence of quinine [39, 53, 70–72]. Therefore, and in order to ensure comparability throughout the study, we used 5 mM quinine test Petri dishes for all experiments, including experiments that only used reward during training. Previous work has demonstrated that the retrieval of fructose reward memories is not influenced by the presence of quinine [53]. To further control whether the test condition does not influence our results, we repeated the complete experiment once more, but with the test performed on a pure, tasteless Petri dish, and obtained very similar results irrespective of the test condition (Fig. S3d-f).

To illustrate how the difference in value between the two substrates determines the preference, we deduced the subjective values of the two rewards directly from the animals’ behaviour during the experiment in Fig. 1a, as the average preference of a group that received AM-reward training minus the average of the baseline preference group (Fig. 1d; for more details, see Materials & Methods). Then, we determined the value difference in each of the experimental groups of Fig. 1c by subtracting the subjective value of the substrate paired with OCT from the value of the substrate paired with AM (Fig. 1e). Finally, we plotted the AM preferences of all groups over the value differences and used a generalized logistic function to describe the relationship between them (Fig. 1f, Fig. S3c,f). Across the data of this study, a generalized logistic function fits the data better than a linear function (Fig. S4).

In summary, we found that larvae learned about the relative value of rewards, and that the value difference between the options during training determined the odour preference in a logistic manner.

### Larvae learn about the relative value of punishments

Next, we asked whether the larvae learn about the absolute or relative value of punishments. To answer this, we performed a similar experiment as shown in Figure 1, but used two different concentrations of quinine as punishment, based on an initial experiment (Fig. S5a-c). We ensured that the larvae can still detect and process the lower of the two quinine concentrations immediately after being exposed to the higher concentration, although they showed a trend of a slightly reduced avoidance (Fig. S5d).

Similar to the previous findings, the animals all displayed the same baseline AM preference after they had been trained with the same quinine concentration in both training dishes, irrespective of the punishment strength (Fig. 2a). When AM was paired with no punishment and thus the absolute punishment value was zero, but the alternative featured different punishment concentrations (Fig. 2b, left), we found that the AM preference was increasing with increasing punishment strength that had been paired with OCT. Finally, when AM had been paired either with the weak punishment (Fig. 2b, middle) or the strong punishment (Fig. 2b, right), in each case the AM preference increased along with the punishment strength that had been paired with OCT. Thus, we concluded that larvae learn about the relative value of a punishment, not the absolute value (Fig. 2c).

**Figure 2.**
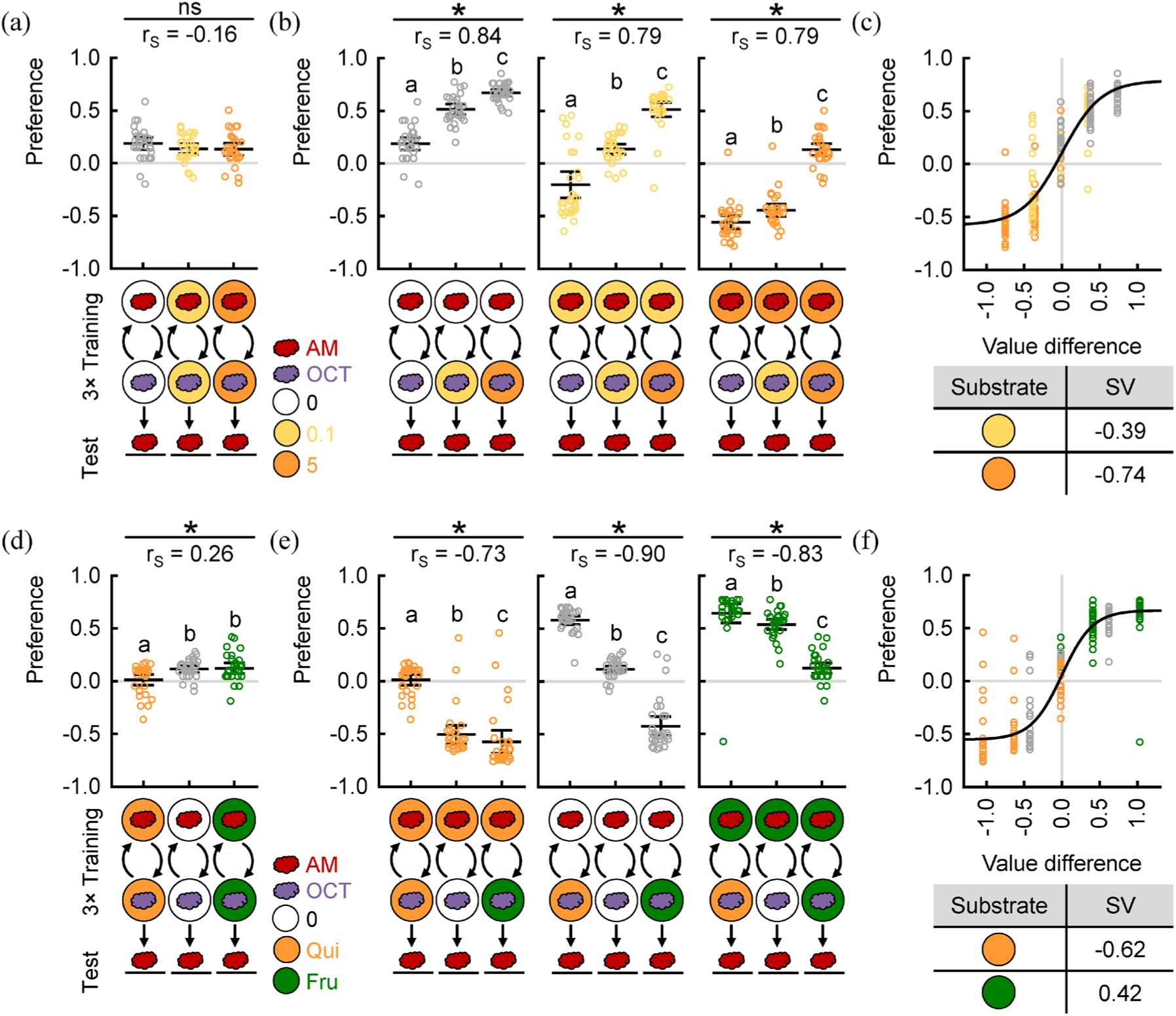
Larvae learn about the relative value of punishments. (a-c) Larvae were trained with two odours, AM and OCT, and different concentrations of quinine (0, 0.1 and 5 mM). (a) The same quinine concentration was presented in both training dishes. (b) The quinine concentration paired with AM was kept constant, but the concentration paired with OCT varied. (c) AM preferences increased with increasing difference in punishment value. The SV of each punishment is displayed below the plot. (d-f) Same as in (a-c), but with combinations of 5 mM quinine as punishment and 2 M fructose as reward. Sample size is 30 each. Note that the groups displayed in (a) and (d) are also included in (b) and (e), respectively. For further details, see Fig. 1. For all raw data along with the detailed results of the statistical tests, see Supplementary table S1.

### Larvae learn about the relative values of combined reward and punishment

After finding that the larvae learned about the relative value of rewards and punishments in separate experiments, we wondered what would happen if we combined reward and punishment in the same experimental setup. When both odours had been paired with the same reward or punishment, we saw a slight but significant difference in AM preference, mostly due to a decreased AM preference after training with quinine (Fig. 2d). We note that this group is an exact repetition of the one in Fig. 2a (right-most group) in which we saw no reduced preference. This difference may point to the level of behavioural variability in our experiments and cautions against over-interpreting small correlations, even if they are statistically significant. When keeping the reward or punishment paired with AM constant, we found, once more, a clear change in AM preference along with the substrate that had been paired with OCT, fitting to the hypothesis of relative value learning (Fig. 2e). We note that pairing one odour with reward, the other one with punishment led to stronger negative or positive preferences than training with only the reward or only the punishment. Thus, we again concluded that larvae learn about the relative values of reward and punishment, not the absolute values. In this experiment, as the value differences between the substrates were particularly large, the logistic relationship (as compared to a simple linear one) between the value difference and the odour preference was particularly striking (Fig. 2f).

### Larvae can learn about relative reward values with only one odour

In addition to the differential, two-odour training paradigm, a one-odour version is used regularly [62, 73–75]. We next asked whether the larvae would also learn the relative reward value in such a paradigm, or whether the presence of two odours during the training was required. We repeated the experiment of Fig. 1 but trained the larvae by pairing the various substrates with only one odour, either AM (Fig. 3a-c) or OCT (Fig. 3d-f).

**Figure 3.**
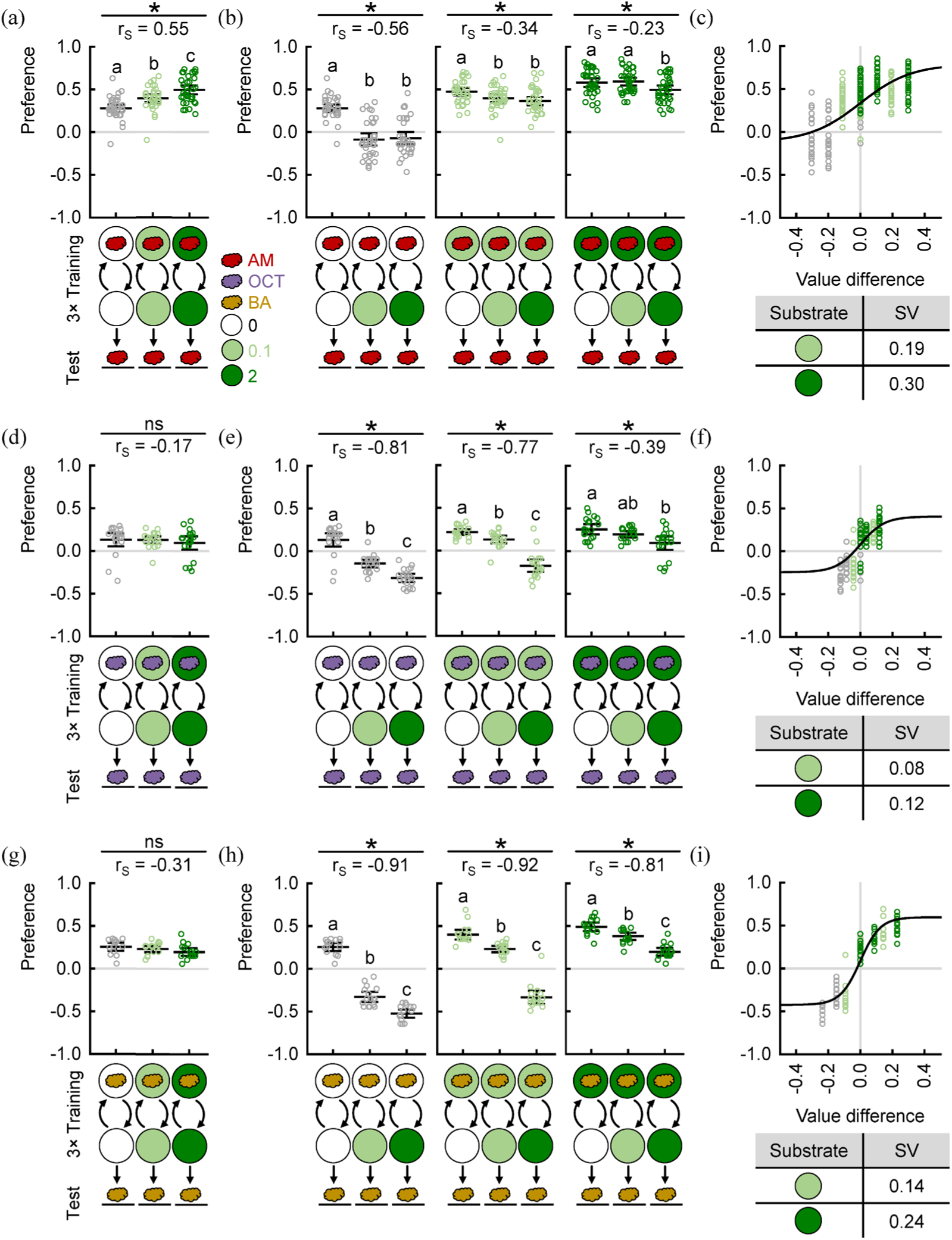
Larvae can learn about the relative value of rewards with one-odour training. (a-c) Larvae were trained only with AM. All other experimental details as in Fig. 1. (a) Larvae were trained with the same fructose concentration in both training dishes. (b) Larvae were trained with the fructose concentration paired with AM kept constant, but the concentration unpaired with AM varied. (c) AM preferences increased with increasing difference in reward value. The SV of each reward is displayed below the plot. (d-f) Same as in (a-c), but for OCT as odour. (g-i) Same as in (a-c), but for BA as odour. Sample size is (a-c) 36, (d-f) 20, (g-i) 16 each. Note that the groups displayed in (a), (d) and (g) are also included in (b), (e) and (h), respectively. For further details, see Fig. 1. For all raw data along with the detailed results of the statistical tests, see Supplementary table S1.

When we trained the larvae with the same fructose concentration in both training dishes, they showed increased AM preferences with increased reward strength which indicates that they learned about the absolute value of the reward (Fig. 3a). Next, we trained the animals such that AM was paired with no reward and thus the absolute reward value was zero, but the alternative featured different reward strengths (Fig. 3b, left). We observed that the AM preference was lower when the alternative substrate had been fructose, suggesting that also the relative value had been learned. When we paired AM with reward, we observed a similar tendency, but dampened the stronger the reward paired with AM had been (Fig. 3b, middle, right).

We repeated the same experiment using OCT (Fig. 3d-f). Here, the OCT preference did not change when the same substrate was used in both training dishes (Fig. 3d), and when the fructose concentration paired with OCT was kept constant, the OCT preference was consistently decreasing with increasing fructose concentrations not paired with OCT (Fig. 3e-f). These results very much matched those from the two-odour experiments and clearly indicated that the larvae learned about the relative but not the absolute value of the rewards.

Finally, we decided to try benzaldehyde (BA), another odour that is often used in larval learning experiments [69, 76], and found results that closely matched those seen with OCT and indicated relative value learning (Fig. 3g-i).

In summary, we found that the larvae can also learn about the relative reward value with only one odour during training. However, the extent to which the relative but not the absolute reward value was learned depended on the trained odour and the reward strength paired with it.

### Larvae learn relatively independent of the training sequence

Finally, we asked whether relative value learned depended on the training sequence which determines the information available to the animal at each step: if AM was presented in the second training phase, the animals could have a memory about the reward strength of the previous phase, and associate the relative increase or decrease with AM. In contrast, if AM was presented first, this information would be not available at the time of AM presentation.

All experiments so far included three training cycles that followed each other immediately and thus do not allow for a precise control of the information available at each step (Fig. S1). Therefore, we switched to a one-trial training paradigm [77]. The training was performed as described for Fig. 1, but the larvae were exposed to each odour-substrate combination only once before they were tested for their AM preference. We first trained the animals such that OCT was always presented first and AM second (Fig. 4a-c). The results were very similar to the three-trial experiments and clearly indicated relative value learning. Then, we trained the animals with the other sequence, presenting AM first and OCT second (Fig. 4d-f). When both odours had been paired with the same reward strength, we found a slight but significant increase of the AM preference with increasing reward strength (Fig. 4d). Surprisingly, when we kept the substrate paired with AM constant, the AM preference decreased with increasing reward strength paired with OCT, indicating relative value learning (Fig. 4e). Overall, the results were very similar between both training sequences. As an additional control, we tested whether the exposure to sugar rewards alone may affect odour preferences in a way that can lead to the observed results (Fig. 4d-f). We found that exposing the animals to reward in absence of any odour decreased the subsequent preference for AM (Fig. S6a). In addition, the sequence of reward exposures mattered: increasing reward strengths from the first to the second dish decreased the subsequent AM preference, decreasing reward strengths increased it (Fig. S6b-c). This trend qualitatively matched the results of the previous experiment, but to a much smaller extent (S6c). Taken together, we concluded that the exposure to sugar alone cannot explain our odour-sugar training results and that, indeed, larvae can learn about the relative reward value irrespective of the training sequence.

**Figure 4.**
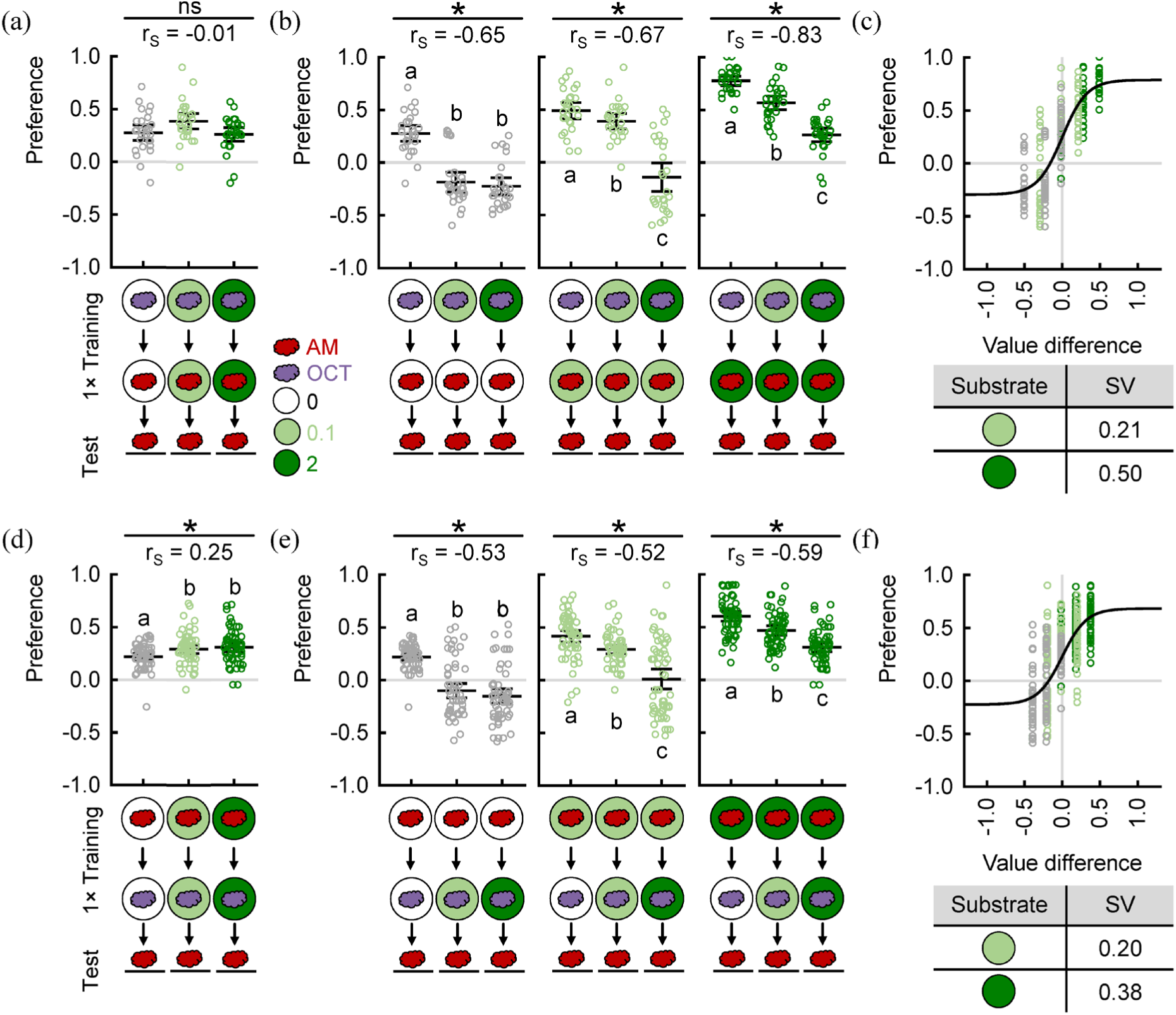
Larvae can learn the relative value of rewards independent of training sequence. Larvae were trained with two odours, AM and OCT, and different concentrations of fructose. Only a single training trial was performed, with (a-c) OCT or (d-f) AM being presented first. (a, d) The same fructose concentration was presented in both training dishes. (b, e) The fructose concentration paired with AM was kept constant, but the concentration paired with OCT varied. (c, f) AM preferences increased with increasing difference in reward value. The SV of each reward is displayed below the plot. Sample size is 28 each for (a-c) and 58 each for (d-f). Note that the groups displayed in (a) and (d) are also included in (b) and (e), respectively. For further details, see Fig. 1. For all raw data along with the detailed results of the statistical tests, see Supplementary table S1.

## Discussion

In this study, we asked whether *D. melanogaster* larvae learn about the absolute or relative values of rewards and punishments during classical conditioning. We designed an experiment that allowed us to distinguish the two hypotheses by pairing each of two odours with rewards or punishments of different strength, followed by testing the preference of one odour alone to assess what the animals had associated with it. Our results clearly match the hypothesis that the values of both training situations were compared and the relative value was associated with one odour, both for rewards (Fig. 1) and punishments (Fig. 2), irrespective of which odour was tested (Fig. S3) as well as of the number and sequence of training trials (Fig. 4). Can these results explained by processes that do not require the comparison of the relative values?

Previous exposure of sugar rewards may change the animals’ motivation and evaluation of further sugar rewards. Indeed, we observed that the larvae showed a reduced preference towards low-sugar when they had experienced the high concentration before (Fig. S2d). This may reduce the subjective reward value that is associated with an odour in the according learning condition and thus explain the differences between training regimen we observed. Alternatively, the learning processes in the two training phases may disrupt each other, a phenomenon called memory interference in Psychology [78–79]. Theoretically, the memory formed about sugar in the first training phase may inhibit learning about the odour-sugar association in the second phase [80]. Likewise, learning about the sugar in the second phase may impair the retrieval of the memory from the first phase [81]. Both processes could lead to differences between our training regimen. However, neither a reduced motivation nor impaired learning or retrieval capabilities can explain an inversion of memory valence – that is, that the animals form an aversive memory about the odour paired with the weaker of two rewards, as we observed it repeatedly (Fig. 1c, 3e, 3h, 4b, S3b, S3e; 6^th^ group from the left).

In contrast, this result is in line with one of the experimental hallmarks of relative value learning in humans: that it can lead to objectively suboptimal decisions. Humans tend to choose options that had been better than alternatives in previous experiences over options that predict a higher absolute value [8, 14–15]. The larvae avoided an odour that had been paired with a potent reward if the alternative had been even better – although no better option was available at the moment of the choice test. This prevented the larvae to find an otherwise positively rated sugar source. Thus, it seems that the way how brains evaluate rewards or punishments during learning is shared across the animal kingdom from insect larvae to humans.

Notably, our results do not exclude that larvae can learn about the absolute values of rewards. What animals learn depends strongly on the specific training procedure. We presented the two training situations in an alternating fashion, each of them paired with a different odour – this procedure likely facilitated a direct comparison of the reward values. A blocked design, or using only one odour during the training, could potentially favour absolute value learning [12]. Indeed, when training only with AM, we observed partially absolute, partially relative value learning (Fig. 3a-c). When using other odours, however, we found complete relative value learning, matching the results of the differential, two-odour training (Fig. 3d-i). The reason for this discrepancy is not known. In any case, this result shows that relative value learning does not require the presence of two alternating stimuli that are associated with the different rewards.

### Relative value learning is consistent with previous studies

To our knowledge, this is the first demonstration of relative value learning in *D. melanogaster*. Previous studies in both larvae and adult flies had used different punishment strengths during training, each paired with a given odour, and then had tested the animals’ choice between these odours [39–40, 44]. These studies demonstrated that the animals can learn how bad a given punishment is, but they could not distinguish between relative and absolute value learning, as it remains possible that the animals learned the absolute value paired with each odour and then compared during the choice test which of the odours predicted the better outcome. The critical advantage of our design is that we tested only a single odour and therefore could directly assess what the animals had learned about it.

A number of studies used an experimental paradigm similar to ours with two odours during the training but only one of them during the test [64, 82–84]. In addition, a one-odour training version is widely used in larvae [49, 53, 61–62, 72–76, 85–86]. In these studies, a single reward or punishment was paired with one odour, while the other training situation was neutral and therefore had a value of zero. Their results are not conclusive for, yet consistent with the hypothesis of relative value learning.

For example, one-odour experiments with rewards always consisted of two types of training regimen: either the odour was paired with the reward while the other training situation featured no odour on a neutral substrate (so-called paired training), or the odour was paired with the neutral substrate while the reward was presented without an odour (so-called unpaired training). For paired training, both absolute and relative value learning would lead to the same outcome, an increased odour preference compared to the baseline. For unpaired training, in contrast, absolute value learning would lead to a baseline level of odour preference because the odour had not been paired with any reward, whereas relative value learning would lead to a preference lower than the baseline because the odour had been paired with the less rewarding of the two substrates. The latter is precisely what had been found in several previous studies and was confirmed in this study (Fig. 3) [61–62, 64, 72]. Analogous results have been observed when using punishments instead of rewards [64, 72] as well when using a training with two odours (Fig. 1c, 2b) [64]. Using a different experimental design, similar results have been found in adults, too. Here, the resulting memories were called “CS+” and “CS-” memories [83].

These observations gave rise to the proposal of two separate learning processes after paired and unpaired training, leading to memories of opposite valences and with separate neurobiological mechanisms [62, 64, 72, 83, 87–88]. However, relative value learning provides a unified framework to explain the results of both kinds of training: the memory’s valence depends on whether the trained odour had been paired with the better or worse of the training situations. This new perspective should be incorporated into the current theoretical, computational and neuronal working models of associative learning in *D. melanogaster*. In the following, we will briefly discuss one plausible neuronal working model.

### A potential neuronal circuit for relative value learning

In *D. melanogaster*, olfactory associative memories are established in the mushroom body, a major insect memory centre (Fig. 5a) [38, 89]. In brief, odours and other stimuli are coded by specific combinations of Kenyon cells (KCs), the intrinsic neurons of the mushroom body. The KCs are intersected by (mostly) dopaminergic modulatory input neurons (DANs) and mushroom body output neurons (MBONs). The innervations of DANs and MBONs compartmentalize the mushroom body into microcircuits between DANs, KCs and MBONs that act as functional units. As a rule, DANs conveying reward signals pair up with avoidance-promoting MBONs, and DANs conveying punishment signals pair up with approach-promoting MBONs. DAN signals are received by the axonal terminals of all KCs and modulate the KC-MBON synapses. The synaptic weights of the KC-MBON synapses thus contain the associative strengths of all stimuli coded by the KCs. Crucially, studies in adults found that the same DAN signal can have opposite effects on internal KC signalling when a KC is active versus inactive [90], and that the KC-MBON synapses can be depressed or potentiated depending on the training contingency [91].

**Figure 5.**
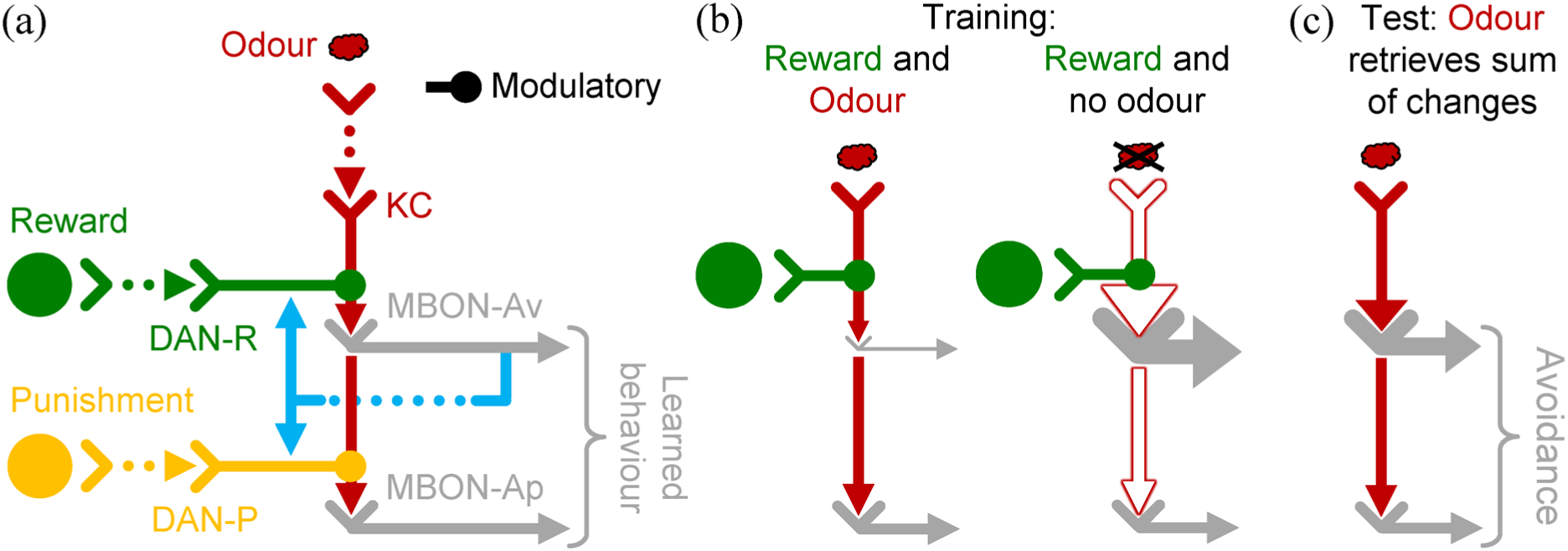
Simplified potential circuitry underlying relative value learning. (a) Mushroom body circuit for associative learning. DAN: dopaminergic neuron; KC: Kenyon cell; MBON: mushroom body output neuron. Two sample compartments are shown, one receiving reward, the other one punishment input. The feedback from MBONs to DANs (blue, only displayed for one compartment) can be direct or indirect and excitatory or inhibitory. Based on [49, 74, 85]. (b) Simple working model of absolute and relative value learning. Left: coincidence of odour and reward depresses the synapses of odour-activated KCs and an avoidance-promoting MBON. Right: a reward received by an inactive KC potentiates the KC-MBON synapses. (c) The KC-MBON synapses integrate the modulations by previous rewards in presence and absence of the odour. In the shown example, the net effect is a potentiated synapse and thus odour avoidance.

The MBONs gather input from all KCs and on the one hand convey this information towards motor control, tilting the behaviour of the animals towards approach or avoidance, depending on the memory [86, 92–94]. On the other hand, they feed this information back either directly, or via one and two step feedback relays to the DANs [74]. The DANs that receive the feedback may innervate the same or other compartments and may even code for the opposite valence [74].

The predominant working model for associative learning in *D. melanogaster* proposes that when a DAN reward signal is received by a KC while it is active (i.e. while the respective odour is present), its output synapses are depressed to an extent matching the reward strength (Fig. 5b, left). This reduces the activity of the avoidance-promoting MBON while leaving the activity of approach-promoting MBONs unchanged and thus shifts the mushroom body’s net output towards approach [86, 92–94]. This process can be seen as the basis for absolute value learning: an odour is associated with the reward strength it coincides with [44].

For relative value learning, the crucial question is how the reward strengths can be compared and the relative value can be computed. In mammals, DANs have been found to compute an associative prediction error that can be interpreted as the difference between the current and previously learned reward strengths and thus as a computation of relative reward value [95–97]. Many computational models assume a similar mechanism in *D. melanogaster*, driven via the feedback from the MBONs to the DANs, but hard evidence for such a mechanism is lacking [74, 98–99].

We propose that the computation of the relative value can occur in the KCs themselves: when a DAN signal is received by an inactive KC (i.e. the respective odour is not present), its output synapses are potentiated to an extent matching the reward strength, leading to a shift in the mushroom body’s net output towards avoidance (Fig. 5b, right) [44, 64, 91]. If rewards of different strengths are presented in presence and absence of an odour, like in our experiment, the final strength of the KC-MBON synapses reflects an integration, in the simplest case a sum, of all modulations during training (Fig. 5c). In other words, the relative reward value is computed within the KC-MBON synapse, as an integration of all positive and negative molecular modulations of the synapse strength.

Interestingly, the proposed model predicts that KC-MBON synapses get potentiated whenever a reward is encountered while a KC is not active. This should decrease the preference for any odour that was not present at that moment. Indeed, we found that exposing the animals to sugar reward alone decreases preferences to a novel odour, although the effect was quite small (Fig. S6a). We also found that the presence of a second training odour was not crucial for relative value learning – what mattered was the difference in reward strength in presence and absence of the to-be-tested odour (Fig. 1, Fig. 3).

Although the proposed working model is very simple, it can be easily expanded to account for additional considerations. For example, humans have been found to compare reward strengths within each learning session [8], and mammals often learn in a context-dependent manner [100–102]. Possibly, also the KCs that get potentiated are limited, e.g. to those that code for stimuli that had been encountered recently, or in the same experimental context. Such a mechanism would prevent an unnecessary potentiation of all KC-MBON synapses whenever a reward is encountered. In adult flies, reward-signalling DANs have been found to be required for memories about the punishment-unpaired odour, but not about the punishment-paired odour [83]. Another study found a MBON pathway that encodes a “better-than-before” signal and activates a reward-signalling dopaminergic pathway, leading to a secondary reward memory in a different mushroom body compartment [44]. In other words, when an odour was paired with the weaker of two rewards or punishments, the modulations of the KC-MBON synapses we propose here may, as a second step, establish a memory of the opposite valence via the opposite-valence DANs. Future research will elucidate these and other pending questions regarding the detailed mechanism of how brains integrate the relative values of options into associative learning, in larvae as well as in other animals. The *D. melanogaster* larva, with its small brain and excellent tools for transgenic manipulations, is perfectly suited to contribute to this endeavour.

## Acknowledgements

We thank A. Fukushima, B. Gerber, G. Jocham, M. Mizunami, P. Tobler and M.L.M. Wang for fruitful discussions and N. Toshima for helpful comments on the manuscript. We are grateful to the Ministry of Education, Culture, Sports and Technology (MEXT), Government of Japan, for financial support to S. Rahman through the MEXT Scholarship.

## Supplementary figures

**Figure S1.**
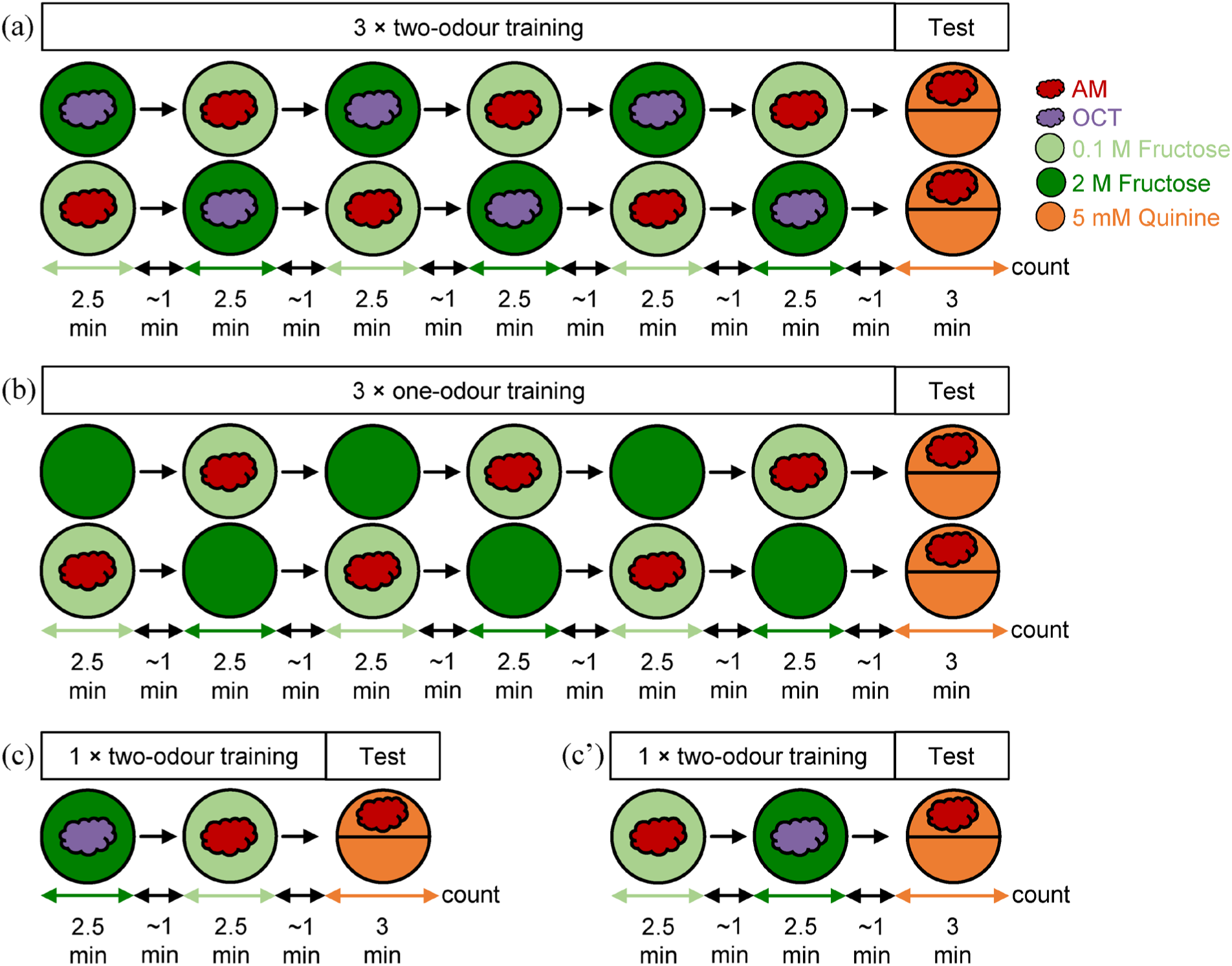
Training procedures in this study. (a) Larvae were trained for 3 cycles with two odours, AM and OCT and different concentrations of fructose. Depicted is group 6 in Table 1 as example, exactly as it was performed for Fig. 1. The larvae spent 2.5 minutes on each dish and were transferred manually with a brush between dishes, which took about 1 minute. After 3 complete cycles, the animals were transferred to a test dish covered with quinine and with only one odour on one side, and after 3 minutes the number of larvae on either side was counted. Each displayed training sequence was used in half of the repetitions of the experiment (samples). For Fig. S3a-c, the identities of AM and OCT were swapped; for Fig. S3d-f, the test dish contained pure agar instead of quinine; for Fig. 2, different quinine concentrations were used during training instead of fructose. (b) One-odour version of the experiment, as it was performed for Fig. 3a-c and S6. OCT was replaced by empty containers (not shown). For Fig. 3d-i, AM was replaced by OCT or BA in training and test. (c) One-trial version of the experiment as it was performed for Fig. 4a-c. (c’) One-trial version of the experiment as it was performed for Fig. 4d-f.

**Figure S2.**
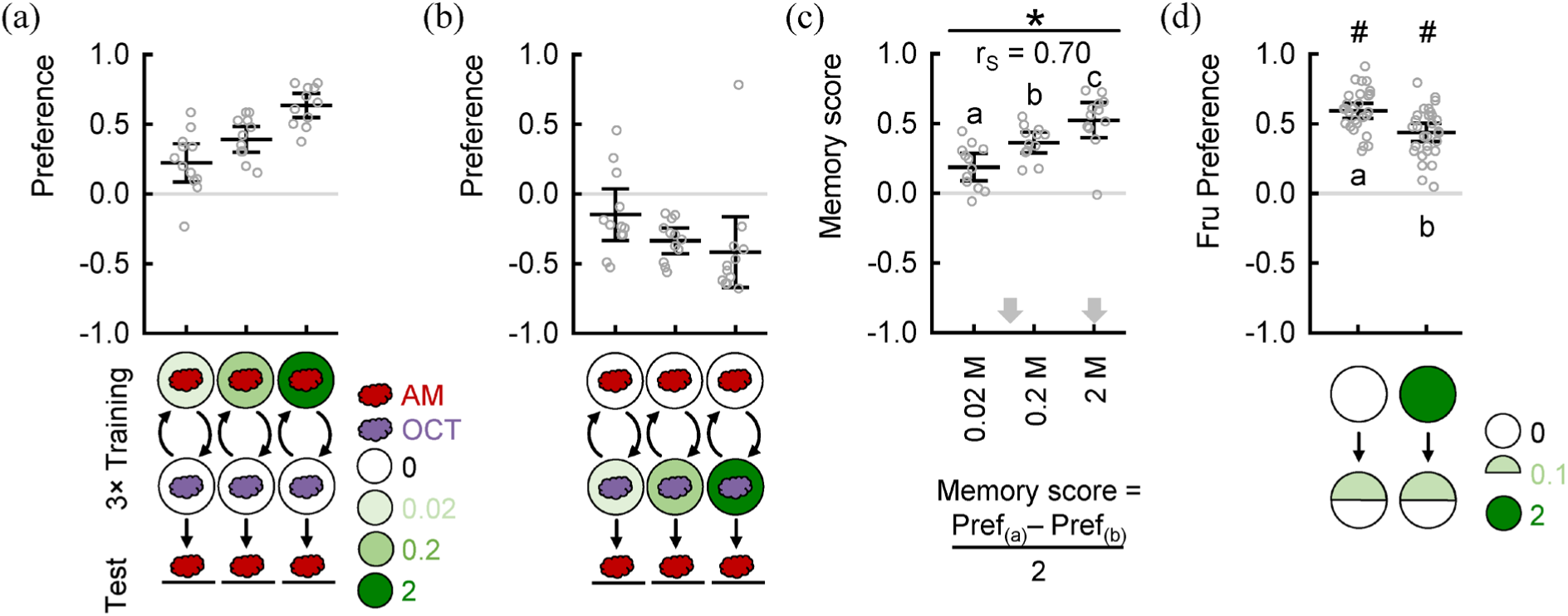
Reward strength increases with increasing fructose concentration. Larvae were trained with two odours, AM and OCT, and different concentrations of fructose (0.02, 0.2 and 2 M). (a) When AM had been paired with the fructose reward, the AM preference increased with increasing fructose concentration. (b) When OCT had been paired with the fructose reward, the AM preference decreased with increasing fructose concentration. (c) Memory scores were calculated as the difference in preference after both kinds of training. They significantly increased with increasing fructose concentrations. The arrows indicate the concentrations picked for the experiments in Figures 1, 3 and S3. (d) Larvae were exposed to either pure agar or agar mixed with 2 M fructose for 2.5 minutes. Subsequently, they were tested for their preference of a 0.1 M fructose substrate in a split Petri dish assay. In both cases, larvae preferred the fructose half. This demonstrates that the sensation and preference towards a low concentration of reward is intact after exposure to a high concentration of reward, as it is used in the learning experiments. Sample size is 12 each for (a-c), and 30 each for (d). Raw data are displayed with the mean ± 95 % confidence interval. In (c), * indicates a significant Spearman Rank test and r_S_ indicates Spearman’s correlation coefficient. In (d), # indicates a significant Wilcoxon Signed Rank test against chance level (0). Different small letters above or below each group indicate significant pairwise difference from each other (Mann-Whitney U tests). For all raw data along with the detailed results of the statistical tests, see supplementary table S1.

**Figure S3.**
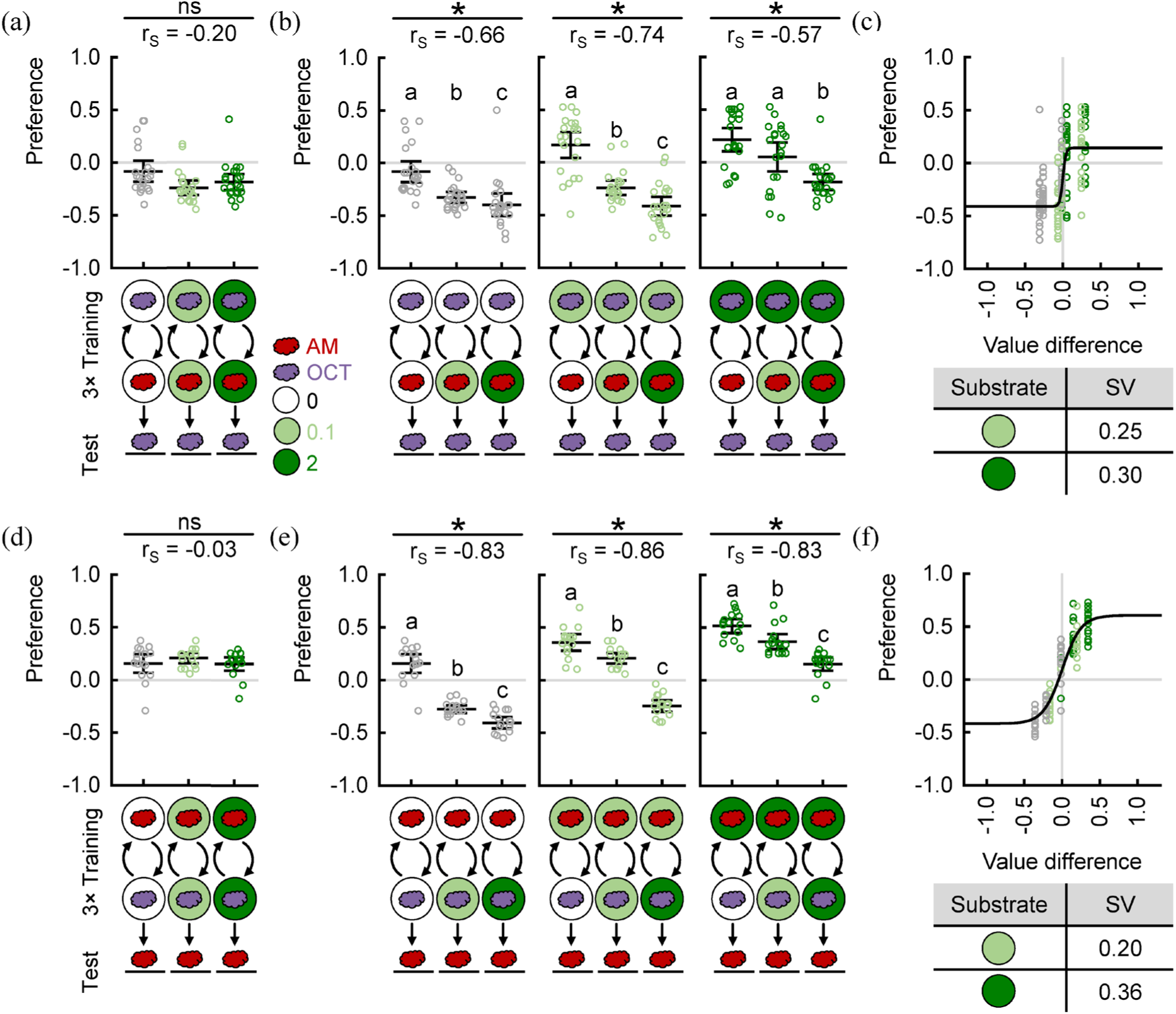
Larvae can learn the relative value of rewards independent of the tested odour or the test condition. (a-c) Larvae were trained with two odours, AM and OCT, and different concentrations of fructose. Afterwards, they were tested for their preference for OCT. (a) When the same fructose concentration was presented in both training dishes, the OCT preference was independent of the fructose concentration. (b) When the fructose concentration paired with OCT was kept constant, the OCT preference decreased with increasing fructose concentration paired with AM, and thus with increasing difference between the two rewards. (c) OCT preferences increased with increasing difference in reward value. The subjective values of each reward, estimated from the animals’ behaviour, are displayed below the plot. (d-f) Same as (a-c), but the animals were tested for their AM preference and on a pure agar, tasteless Petri dish, in contrast to all other experiments that used a quinine test dish. The results were very similar to (a-c) and to Figure 1. Sample size is 22 each for (a-c), and 16 each for (d-f). Note that the groups displayed in (a) and (d) are also included in (b) and (e), respectively. Raw data are displayed with the mean ± 95 % confidence interval. * indicates a significant Spearman Rank test and r_S_ indicates Spearman’s correlation coefficient. Different small letters above each group indicate significant pairwise difference from each other (Mann-Whitney U tests). For all raw data along with the detailed results of the statistical tests, see supplementary table S1.

**Figure S4.**
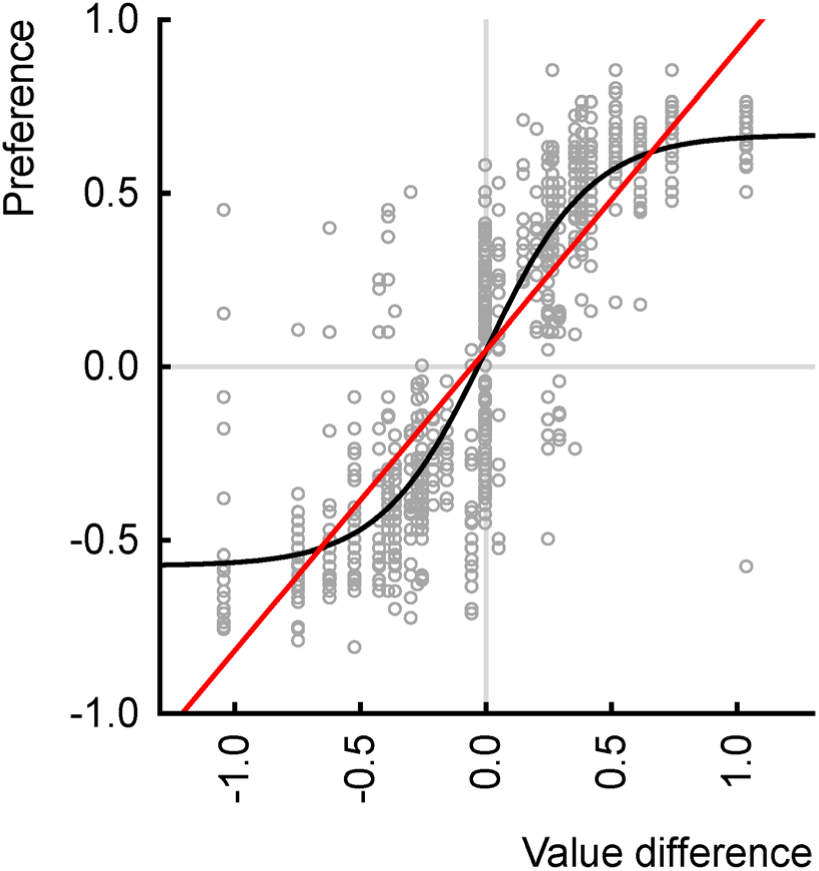
The relationship between the value difference and the odour preference is logistic. This graph displays all data of this study using three training trials with two odours (Fig. 1, 2, S3). To compare a generalized logistic function with a simple linear function, we calculated the Akaike information criterion (AIC) that is regularly used to assess the goodness of a fitting. The AIC for the generalized logistic model was lower than the AIC for the linear function (−390.1 versus −133.9), indicating that the generalized logistic model fitted the data better than the linear one.

**Figure S5.**
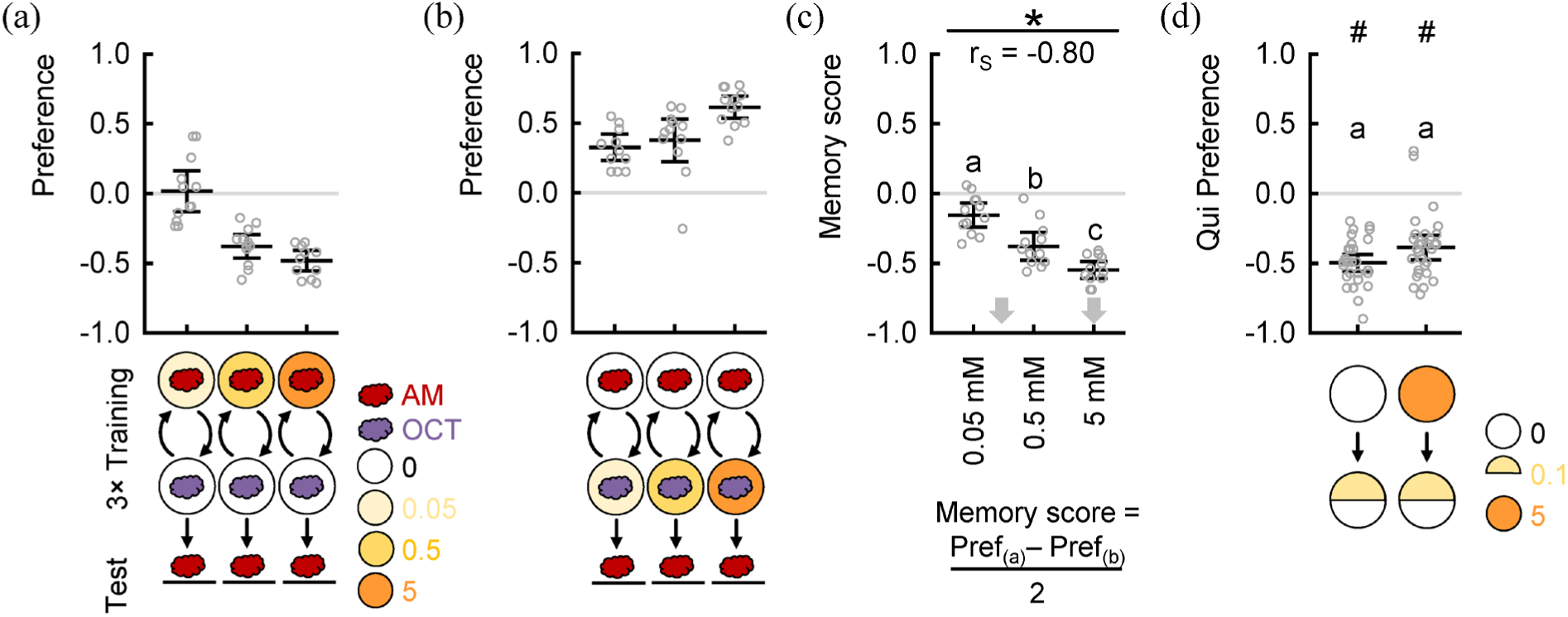
Punishment strength increases with increasing quinine concentration. Larvae were trained with two odours, AM and OCT, and different concentrations of quinine (0.05, 0.5 and 5 mM). (a) When AM had been paired with the quinine punishment, the AM preference decreased with increasing quinine concentration. (b) When OCT had been paired with the quinine punishment, the AM preference increased with increasing quinine concentration. (c) Memory scores were calculated as the difference in preference after both kinds of training. They got significantly more negative with increasing quinine concentrations. The arrows indicate the concentrations picked for the experiments in Figure 2. (d) Larvae were exposed to either pure agar or agar mixed with 5 mM quinine for 2.5 minutes. Subsequently, they were tested for their preference of a 0.1 mM quinine substrate in a split Petri dish assay. In both cases, larvae avoided the quinine half. This demonstrates that the sensation and repulsion towards a low concentration of punishment is intact after exposure to a high concentration of punishment, as it is used in the learning experiments. Sample size is 12 each for (a-c), and 30 each for (d). Raw data are displayed with the mean ± 95 % confidence interval. In (c), * indicates a significant Spearman Rank test and r_S_ indicates Spearman’s correlation coefficient. In (d), # indicates a significant Wilcoxon Signed Rank test against chance level (0). Different small letters above each group indicate significant pairwise difference from each other (Mann-Whitney U tests). For all raw data along with the detailed results of the statistical tests, see supplementary table S1.

**Figure S6.**
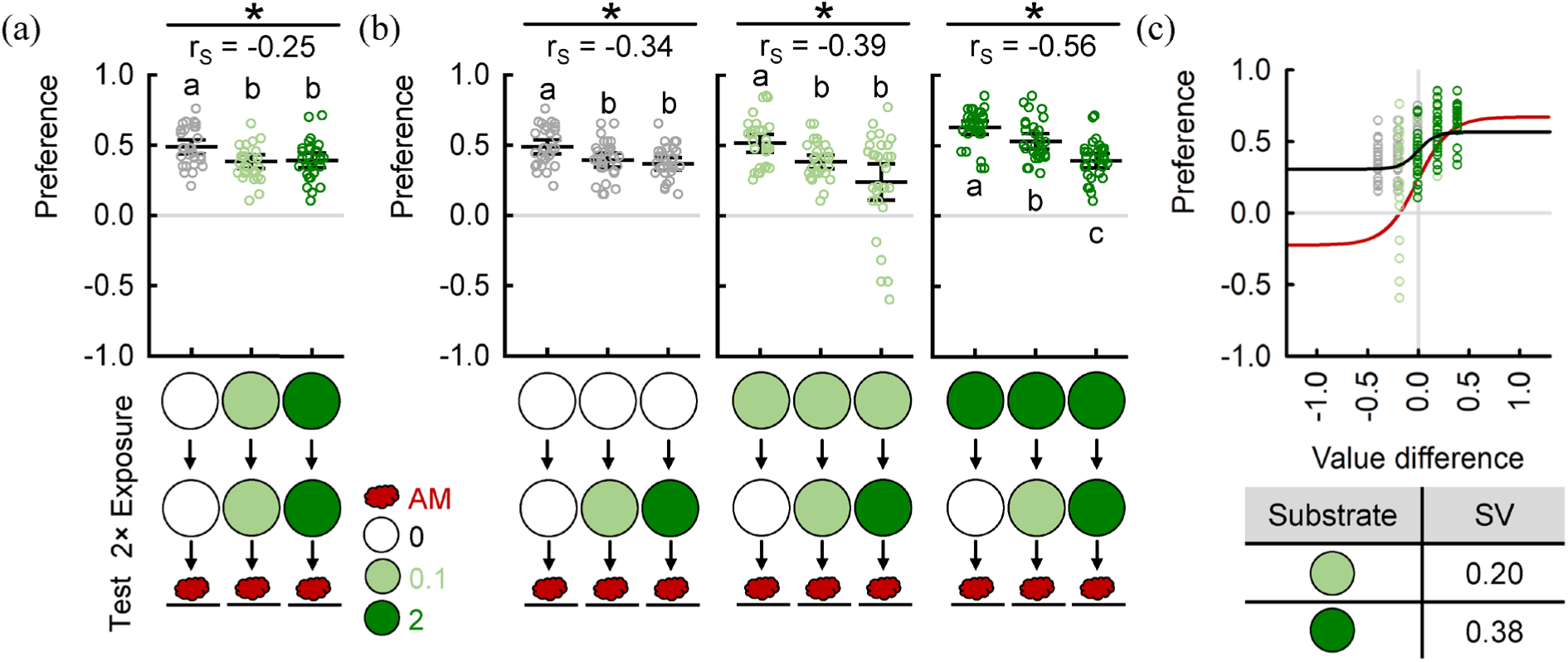
Sugar exposure affects odour preference in a sequence-dependent manner. Larvae were sequentially exposed to two dishes containing various concentrations of fructose. Afterwards, they were tested for their preference for AM. (a) When the same fructose concentration was presented in both exposure dishes, the AM preference was decreasing after fructose exposure relative to the baseline group. (b) When the fructose concentration in the first dish was kept constant, the AM preference decreased with increasing fructose concentration in the second dish. (c) AM preferences increased with increasing difference in reward value, which was calculated in this case as the reward value of the first minus the second exposure dish. The subjective values of each reward were estimated from the animals’ behaviour in the experiment displayed in Fig. 4d-f which had been performed in parallel. In addition to the generalized logistic function fitted to the displayed data (black), the corresponding fit of Fig. 4f is shown (red) for easy comparison. Sample size is 30 each. Note that the groups displayed in (a) are also included in (b). Raw data are displayed with the mean ± 95 % confidence interval. * indicates a significant Spearman Rank test and r_S_ indicates Spearman’s correlation coefficient. Different small letters above or below each group indicate significant pairwise difference from each other (Mann-Whitney U tests). For all raw data along with the detailed results of the statistical tests, see supplementary table S1.

